# Pathology foundation-model embeddings reveal a morphological landscape of breast lesions beyond diagnostic categories

**DOI:** 10.64898/2026.11.02.760689

**Authors:** Yuki Muroyama, Eichi Takaya, Takashi Suzuki

**Author notes:** Corresponding author: Yuki Muroyama, MD, PhD, Address: SiRIUS Institute of Medical Research, Tohoku University, 1-1 Seiryo-machi, Aoba-ku, Sendai, Miyagi, 980-8574, Japan.

## Abstract

**Background:** Despite rapid advances in computational pathology, including foundation-model embeddings, most approaches remain classifier-based. Breast lesion pathology is an area of unmet clinicopathological need and a compelling setting for morphology-based modeling: diagnostic categories occupy overlapping morphological spectra and reflect heterogeneous pathways of tumor development, thereby posing substantial challenges for pathological diagnosis and subsequent clinical decision-making. However, approaches to delineate morphological continua across diagnostic categories and tumor development remain understudied. Here, we leveraged morphology-based foundation-model embeddings to define a morphologic state space across breast lesion categories and dissect how these embeddings could reflect a pathobiological continuum or sequence associated with tumor development.

**Methods:** To model breast lesion categories in continuous morphology space, we established a leakage-controlled, patient-aware workflow using the frozen foundation model Prov-GigaPath and the BReAst Carcinoma Subtyping (BRACS) dataset, a pathologist-annotated hematoxylin-and-eosin (H&E) breast lesion dataset spanning normal tissue, benign proliferative lesions, atypical lesions, ductal carcinoma in situ (DCIS) and invasive carcinoma (IC). Using the BRACS training split, we defined a train-derived principal component analysis (PCA) morphology space. We then performed two complementary analyses: normal-rooted k-nearest-neighbor (kNN)-Dijkstra graph-distance scoring to derive a trajectory-like morphology score without imposing the full ordinal diagnostic sequence; pairwise Energy distance and squared maximum mean discrepancy with a radial-basis-function kernel (RBF-MMD²) to quantify order-free distributional separation among diagnostic categories. Validation and test data were projected into the train-defined reference space to assess reproducibility.

**Results:** Prov-GigaPath embeddings defined a train-derived morphology space with category-associated organization. Without imposing the full diagnostic sequence, normal-rooted kNN-Dijkstra graph distances recovered a trajectory-like morphology arrangement from normal/benign to malignant lesions. Order-free Energy distance and RBF-MMD² further supported graded separation among lesion categories. Validation and test datasets preserved the pathological organization. Furthermore, morphological embeddings dissected DCIS heterogeneity associated with concurrent invasive carcinoma, without requiring additional molecular information.

**Conclusions:** Leveraging morphology-based foundation-model embeddings, we defined a morphologic state space across breast lesion categories, allowing nuanced positioning of breast lesions along a pathobiological continuum associated with tumor development. Our approaches pave the way for extracting biologically and clinically relevant information from routine histopathology to guide molecular testing, individualized risk assessment, and cost-efficient clinicopathological decision-making.

**Highlights:**

- Overlapping morphological features challenge breast diagnostic categorization
- Foundation-model embeddings defined a morphologic state space
- Embeddings allow nuanced positioning along morphological continua across categories
- Embeddings dissected DCIS heterogeneity associated with concurrent invasive carcinoma
- Latent morphological signals in H&E slides may refine individualized risk assessment

## Background

Machine-learning applications in digital pathology are rapidly evolving, and a major focus has been on assigning histology images to predefined diagnostic categories(1, 2). This paradigm has produced powerful diagnostic classifiers. Yet, pathological lesions often exist along a biological and morphological continuum. Tumors and disease processes do not always fall into discrete, sharply demarcated categories; rather, borderline or intermediate lesions frequently exhibit overlapping features and thus cannot be fully captured with conventional classification-based approaches. However, approaches and models that delineate the morphological continuum across diagnostic categories and tumor development have been understudied.

Genomic, clonal and single-cell approaches have provided important biological views of tumor evolution and precursor states, but these assays remain costly, tissue-consuming and incompletely integrated into routine diagnostic workflows(3–5). In contrast, hematoxylin-and-eosin (H&E) staining remains the most widely available, timely, and cost-efficient representation of tissue morphology. In daily practice, pathologists often encounter situations where timely diagnosis based only on morphological features of H&E slides is required, although morphological distinction can be challenging. Prior work has also shown that histological image patterns can encode molecular and genetic information, supporting the broader concept that H&E morphology contains biological signals beyond conventional diagnosis(6). Thus, maximizing and leveraging the information extracted from histopathological morphology are essential not only for biological understanding but time- and cost-efficient clinicopathological decision-making, including the prioritization of molecular testing to inform targeted therapy selection and precision oncology.

Breast cancer is the most common cancer among women worldwide and remains a leading cause of cancer-related mortality(7, 8). Pathological diagnosis is essential to determine the range and scope of therapeutic options, including but not limited to the use and regimen of neoadjuvant chemotherapy (NAC), hormonal therapy, extent of surgical resection, and post-operative adjuvant therapies(9). In breast pathology, diagnostic entities such as benign proliferative lesions, atypical lesions, ductal carcinoma in situ (DCIS) and invasive carcinoma (IC) are defined as discrete categories for clinical practice, yet many of these lesions occupy overlapping morphological and biological spectra, posing a diagnostic challenge. Indeed, recent advances in molecular pathology emerge to reveal the heterogeneity of breast cancer development, premalignant lesions, and pathways to develop invasive carcinoma from DCIS(10–12), further adding complexity. Collectively, these features make breast lesion pathology an area of unmet clinicopathological need and a compelling setting for morphology-based modeling beyond discrete diagnostic labels.

Pathology foundation models now provide a way to represent H&E images as high-dimensional morphology embeddings without training a task-specific classifier for each diagnostic problem(13). We reasoned that such embeddings could be used not only to classify lesions, but also to model lesion categories as distributions in a shared morphology state space. This approach is particularly relevant for breast precursor and atypical lesions, where diagnostic categories are clinically important but may not form a strictly linear biological progression.

To delineate the morphological continuum across breast diagnostic categories associated with tumor development, we used the BReAst Carcinoma Subtyping (BRACS)(14), a pathologist-annotated H&E breast lesion dataset spanning normal tissue, benign proliferative lesions, atypical lesions, ductal carcinoma in situ and invasive carcinoma. BRACS was well suited to this analysis because it includes seven consensus-annotated categories across the breast lesion spectrum, including usual ductal hyperplasia (UDH), flat epithelial atypia (FEA) and atypical ductal hyperplasia (ADH), which are essential for evaluating intermediate morphology states. We extracted frozen Prov-GigaPath(15) embeddings from BRACS regions of interest (ROIs) and analyzed them in a patient-aware, train-defined morphology space. Prov-GigaPath was selected because it is an open-weight pathology foundation model pretrained on large-scale real-world whole-slide pathology data, including 1.3 billion 256 × 256 pathology image tiles from more than 171,000 whole-slide images, making it well suited for tile-level embedding and ROI-level aggregation(15).

In this study, we show that breast histopathology images exhibited reproducible organization within an embedding-defined morphology space, consistent with a pathobiological continuum, without imposing the full diagnostic sequence or using molecular biological information.

## Methods

### Dataset and diagnostic categories

We analyzed hematoxylin-and-eosin-stained breast lesion images from the BRACS cohort(14). Regions of interest were grouped according to the BRACS diagnostic categories: normal tissue (N), pathological benign lesion (PB), usual ductal hyperplasia (UDH), flat epithelial atypia (FEA), atypical ductal hyperplasia (ADH), ductal carcinoma in situ (DCIS), and invasive carcinoma (IC). The analysis was designed to evaluate morphology-associated organization in a frozen histopathology embedding space, rather than to train a diagnostic classifier.

### Patch extraction and quality control

BRACS region of interest (ROI) images were provided at 40× magnification. ROIs were preprocessed and divided into image patches, which underwent tissue and background quality control. Patch-level features were extracted from retained patches. ROIs yielding no retained patches after quality control were not included in embedding generation.

### Prov-GigaPath embedding representation

Frozen Prov-GigaPath embeddings were used as morphology representations(15). Patch-level embeddings were aggregated to ROI-level vectors. The downstream analyses used ROI-level Prov-GigaPath embeddings(15) as the primary input. No classifier was trained on the BRACS labels for the downstream morphology-space analyses. The resulting ROI-level embedding table was used for downstream analyses including dimensionality reduction, graph-based morphology scoring, and distributional distance analysis.

### Patient-aware metadata linkage and split-integrity audit

BRACS metadata were used to link ROI records to whole-slide images (WSIs) and patients and to verify split assignments(14). After generation of the ROI-level embedding table and before principal component analysis (PCA) fitting or any downstream morphology-space analysis, we performed a patient-aware split-integrity audit across the train, validation, and test partitions. To control data leakage(16, 17), one patient, assigned to both the training and validation partitions in the full BRACS WSI manifest, was conservatively excluded, and all available ROI records from this patient were removed under a split-integrity rule applied before downstream analysis. This exclusion was applied before all reported downstream analyses, including PCA fitting, graph construction, distance calculations, and bootstrap analyses.

### Train-defined PCA morphology space

Principal component analysis (PCA) was fitted using only training-set ROIs after patient-level exclusion(18). The learned PCA transform was then applied without refitting to training, validation, and test ROIs to define a common train-derived morphology space. Diagnostic labels were not used to fit PCA. Validation and test ROIs were not used to define PCA axes and explained-variance ratios were calculated from the training set only. Dimensionality sensitivity was assessed using lower- and higher-dimensional representations of the same train-fitted PCA.

### Normal-rooted kNN-Dijkstra graph-distance analysis

To assess graph-based morphology organization without imposing the full ordinal diagnostic sequence, we constructed k-nearest-neighbor (kNN) graphs in the train-defined PCA space using training ROIs only. Within this space, a representative Normal training ROI was selected as the graph root. Dijkstra graph distances from this root were computed and normalized to generate a Normal-rooted graph-distance morphology score(19). Graph parameters were selected using a pre-specified connectivity criterion rather than by maximizing association with diagnostic class order. Validation and test ROIs were projected onto the fixed training reference without being added to the training graph. Projection distances were compared with a training-derived reference threshold to identify out-of-reference projections.

### Pairwise distributional distance analysis

Class-specific embedding distributions were compared using pairwise Energy distance as the primary distributional metric(20). Analyses were performed separately in the training, validation, and test splits using the primary representation of the fixed train-fitted PCA space. Biased V-statistic estimates, which include self-pairs in the within-class terms, were used in the primary figures.

Squared maximum mean discrepancy with a radial-basis-function kernel (RBF-MMD²) was used as a complementary sensitivity analysis(21). RBF bandwidths were defined using training data and applied unchanged to all three splits. Biased MMD² estimates were used in the figures.

Diagnostic labels were used only to define the distributions being compared; no ordinal class sequence or adjacency structure was imposed in calculating either metric.

### Patient-aware bootstrap analysis

To account for within-patient dependence, patient-level resampling variability in pairwise Energy-distance estimates was assessed using patient-level bootstrap resampling(22). Patients were resampled as blocks to preserve within-patient dependence. Bootstrap medians and 95% percentile intervals were calculated for the pairwise Energy-distance estimates shown in the main figure. These intervals were interpreted as summaries of patient-level resampling variability rather than as formal null-hypothesis tests.

### DCIS context analysis

After upstream exclusion, all DCIS ROIs from the training, validation and test splits were analyzed in the fixed train-derived reference space. Same-WSI IC co-observation was defined from the BRACS WSI-level ROI-distribution annotation as the presence of at least one IC-labeled ROI on the same WSI. For the primary independent-patient analysis, five patients contributing DCIS WSIs in both same-WSI contexts were excluded, leaving 26 patients with no observed same-WSI IC and 18 patients with observed same-WSI IC. Normal-rooted kNN-Dijkstra scores were summarized at the patient level, accounting for the WSI structure. Groups were compared using an exact two-sided Mann–Whitney U test. Rank-biserial correlation was reported as an effect-size measure, with positive values indicating higher scores in the observed same-WSI IC group. The difference in group medians was accompanied by a 95% percentile bootstrap confidence interval based on within-group patient-level resampling, and a label-permutation test stratified by training, validation and test split was used as a sensitivity analysis.

Sensitivity analyses used a broader patient-level definition based on whether any IC annotation was observed in the available BRACS material, used the five mixed-context patients for a within-patient paired comparison, repeated the same-WSI analysis using alternative PCA representations, and restricted the comparison to validation/test patients. The influence of same-patient IC nodes on training-set graph scores was assessed by excluding these nodes from the graph reference; validation and test patients were already absent from the training graph.

For descriptive visualization in the train-defined PC1–PC2 plane, DCIS morphology was represented by one WSI-balanced vector per patient, matching the multivariate analysis. All retained ROI coordinates were shown faintly for orientation. The two-dimensional display was not used for statistical inference; primary graph-score and distributional analyses used the primary PCA representation. For multivariate distributional comparisons, one WSI-balanced PC-space vector was used per patient. Biased Energy distance was used as the secondary distributional statistic, with biased RBF-MMD² as a metric sensitivity analysis. The RBF bandwidth was defined using training data without group labels. Significance was assessed using patient-label permutations stratified by data split.

All DCIS analyses were exploratory, and the reported P values were interpreted as nominal rather than confirmatory.

## Results

### Leveraging pathology foundation-model embeddings to model morphologic state distribution space of breast tumor lesions

To model breast lesion categories in morphology space, we established a patient-aware workflow using BRACS ROI embeddings generated with Prov-GigaPath (**Fig. 1**). After patient-level split auditing and exclusion of one overlapping patient, the final analysis cohort contained 4,376 ROIs from 384 WSIs and 150 patients, with no residual split leakage detected at the available patient, WSI or slide levels.

**Fig. 1.**
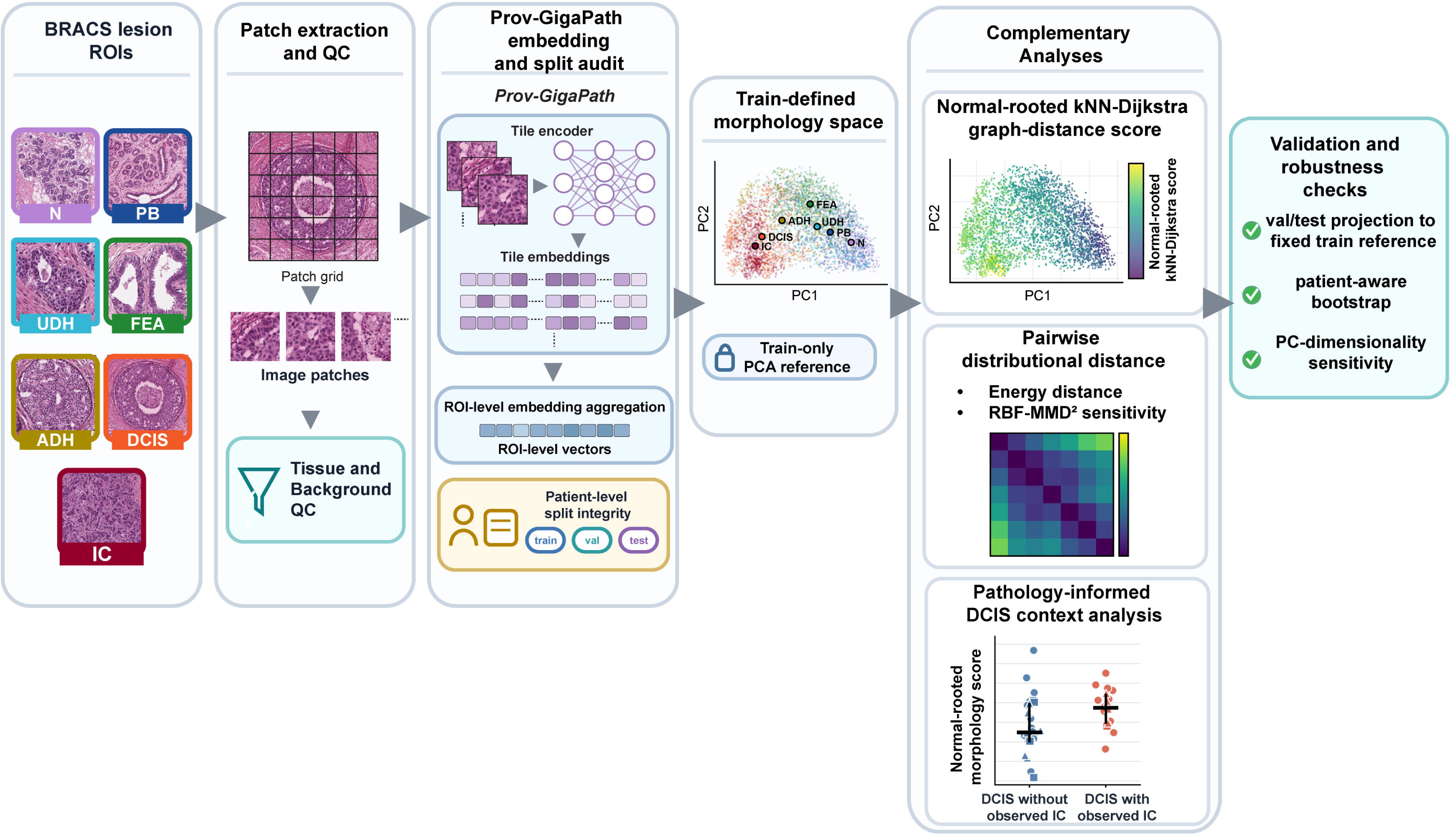
Overview of the patient-aware BRACS morphology-space analysis workflow. BRACS lesion ROIs from normal tissue (N), pathological benign lesion (PB), usual ductal hyperplasia (UDH), flat epithelial atypia (FEA), atypical ductal hyperplasia (ADH), ductal carcinoma in situ (DCIS), and invasive carcinoma (IC) were divided into image patches, subjected to quality control (QC) and tissue/background filtering, and encoded with the frozen Prov-GigaPath tile encoder. Tile-level embeddings were aggregated into ROI-level vectors. Patient metadata were used to audit train, validation, and test partitions; one patient was excluded before all downstream analyses because of train/validation overlap. PCA was fitted only on the leakage-controlled training split. The fixed train-derived reference space supported two complementary analyses: Normal-rooted k-nearest-neighbor (kNN)-Dijkstra graph-distance scoring, and pairwise Energy-distance and RBF-MMD² analyses. Validation and test ROIs were projected onto the fixed training reference, and robustness was evaluated using patient-aware and sampling-sensitivity analyses.

Using the BRACS training split, we defined a train-derived PCA morphology space. Within this fixed reference space, we performed two complementary analyses: Normal-rooted kNN-Dijkstra graph-distance scoring to derive a trajectory-like morphology score without imposing the full ordinal diagnostic sequence; pairwise Energy distance and squared maximum mean discrepancy with a radial-basis-function kernel (RBF-MMD²) to quantify order-free distributional separation among diagnostic categories. Validation and test ROIs were then projected into the train-defined reference space to assess reproducibility and out-of-reference behavior (**Fig. 1**).

Because patient-level leakage can inflate apparent performance in medical image analysis(16, 17), we first linked ROI-level embeddings to BRACS patient metadata and audited the train, validation and test partitions before downstream analyses. This identified one patient-level train/validation overlap in the full WSI manifest, which was removed before downstream analyses including PCA fitting, graph construction, distance calculations, and bootstrap analyses (**Fig. 1**). Thus, the subsequent analyses were performed in a leakage-controlled, patient-aware BRACS ROI embedding cohort.

### Train-defined morphology space reveals BRACS category-associated organization

We first asked whether ROI-level Prov-GigaPath embeddings contained visible morphology organization across BRACS diagnostic categories. PCA was fitted only on training ROIs after patient-level exclusion and then applied without refitting to training, validation, and test ROIs. The first three principal components explained 24.0% of the training-set variance. Across the PC1–PC2 and PC1–PC3 projections and the three-dimensional PC1–PC2–PC3 overview, projected ROIs from all three splits and pooled ROI-level class centroids showed broad category-associated organization, with N and PB concentrated toward one side of the train-defined space and DCIS and IC toward the opposite side, whereas UDH, FEA and ADH occupied partially overlapping regions between these extremes (**Fig. 2A–C**).

**Fig. 2.**
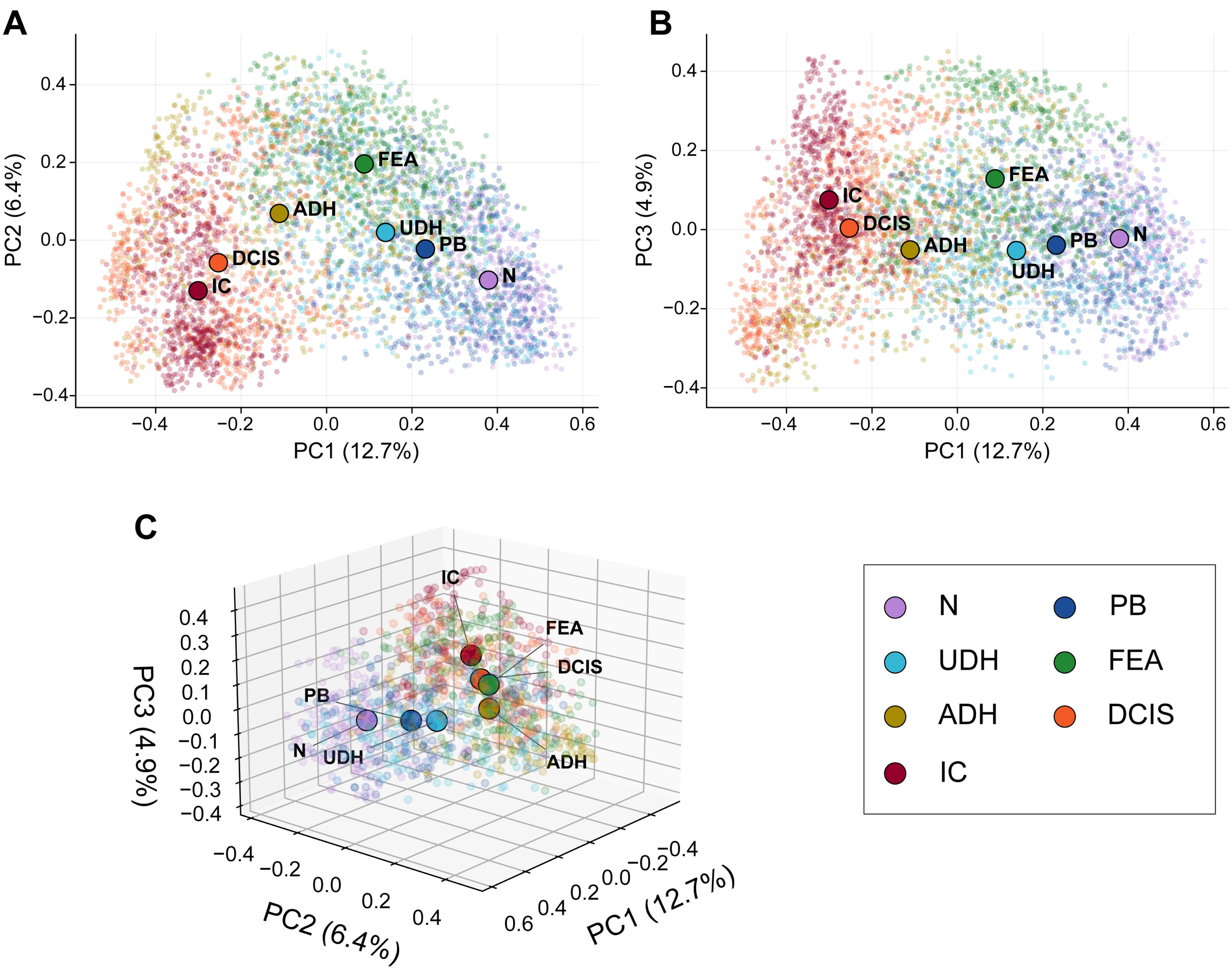
Train-defined PCA morphology-space geometry across BRACS lesion categories. **(A)** ROI-level distribution in the PC1-PC2 plane, colored by pathology class, with class centroids overlaid and labeled. **(B)** ROI-level distribution in the PC1-PC3 plane. **(C)** Three-dimensional PCA overview using PC1, PC2, and PC3, with a class-balanced subsample of ROI points and class centroids. PCA was fitted exclusively on leakage-controlled training ROIs and the fixed transform was applied to validation and test ROIs.

The principal graph-based and distributional patterns were retained when the analyses were repeated using alternative PCA representations under the same parameter-selection rules, indicating that the downstream organization was not specific to the primary representation.

Collectively, these data indicate that frozen Prov-GigaPath embeddings define a train-derived morphology space with visible BRACS category-associated structure but substantial overlap among categories.

### Normal-rooted kNN-Dijkstra graph distances recover trajectory-like morphology organization

We next tested whether local neighborhood structure in morphology embedding space could define a Normal-rooted morphology score without imposing the full ordinal diagnostic sequence. We constructed k-nearest-neighbor graphs using training ROIs only and used a representative Normal training ROI as the root. Conceptually, the graph-based score reflects how far a training ROI is from the Normal root along a network connecting nearby ROIs in embedding space.

Candidate k values were evaluated for connectivity to the Normal root, and the primary k was selected by a pre-specified graph criterion rather than by maximizing concordance with diagnostic class order. The resulting morphology scores showed stable class-level organization across alternative graph settings.

To visualize this graph-derived score in the train-defined PCA projection, we displayed training ROIs in the same train-fitted PC1-PC2 coordinate frame used in Fig. 2A and colored them by the Normal-rooted kNN-Dijkstra morphology score. The score was computed on the kNN graph constructed in the primary PCA space, whereas the PC1-PC2 plane was used only for visualization. The visualization showed a broad increase in the Normal-rooted kNN-Dijkstra morphology score from N-associated regions toward IC-associated regions (**Fig. 3A**).

**Fig. 3.**
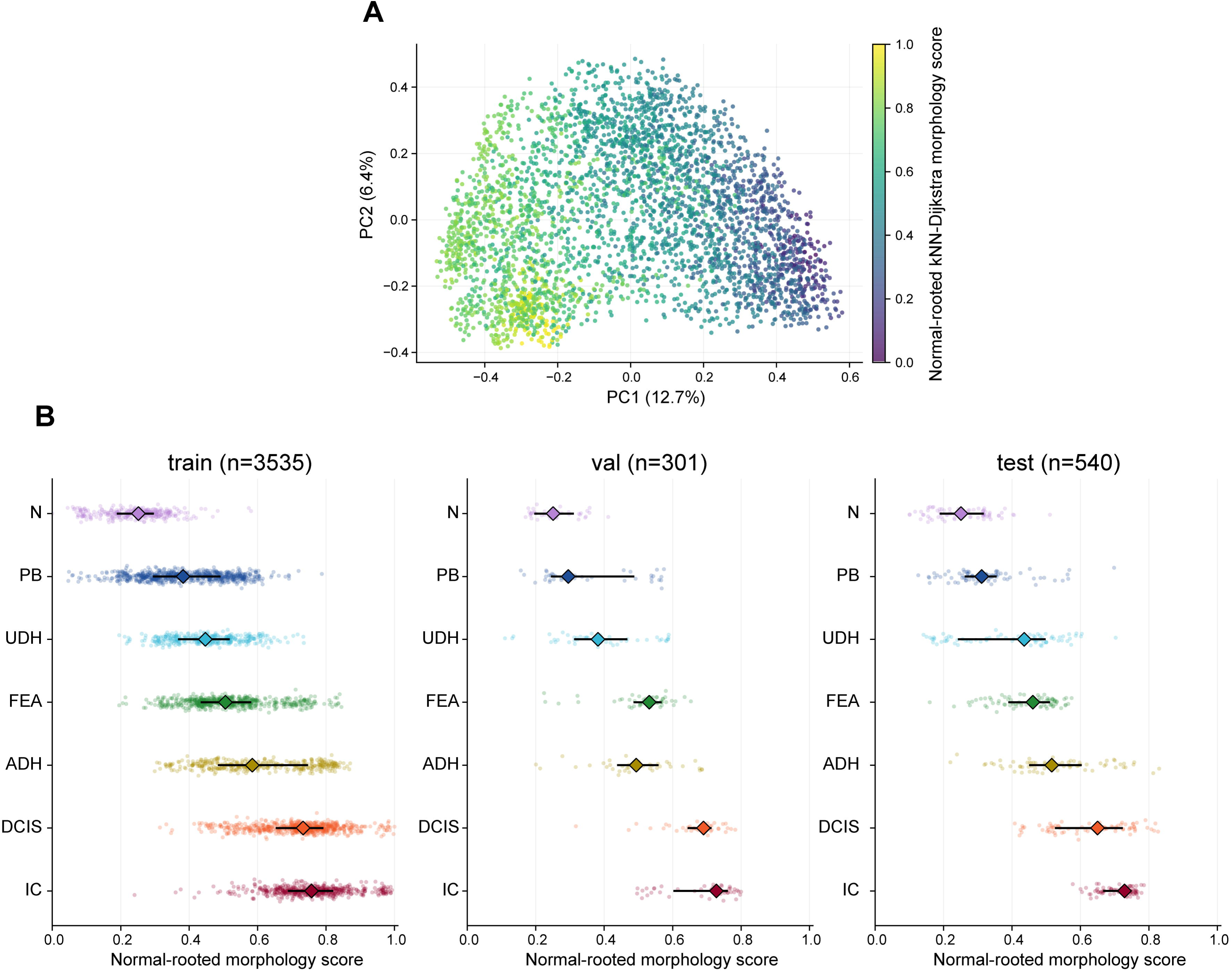
Normal-rooted kNN-Dijkstra graph distances define a reproducible morphology score. **(A)** Training ROIs displayed in the same train-fitted PC1-PC2 coordinate frame and plotting geometry used in Fig. 2A and colored by the graph-distance score. The score was computed on the kNN graph constructed in the primary PCA space. **(B)** ROI-level Normal-rooted embedding-space graph distance score distributions in the training, validation, and test splits. Validation and test ROIs were projected onto the fixed training reference. Points are individual ROIs, diamonds indicate class medians, and black bars indicate interquartile ranges.

At the primary k, ROI-level morphology scores preserved a reproducible class-associated organization across the training, validation, and test splits, with substantial within-class heterogeneity (**Fig. 3B**). In the training split, class medians increased in the order N, PB, UDH, FEA, ADH, DCIS, and IC, although this full sequence was not used to compute the score.

Validation and test projections retained strong rank associations with the diagnostic-category index (Spearman’s ρ = 0.779 and 0.815, respectively), with a local FEA/ADH interchange in validation and closely positioned UDH and FEA medians in test data (**Fig. 3B**). Median nearest-training-reference distances were below the training-derived reference threshold in both splits, whereas 34.9% of validation ROIs and 26.7% of test ROIs exceeded this threshold. Restriction to in-reference ROIs retained strong rank associations (Spearman’s ρ = 0.771 in validation and 0.791 in test data), indicating that the broad rank structure was not driven solely by out-of-reference projections.

Together, these data indicate that Normal-rooted kNN-Dijkstra graph distances recover a reproducible morphology-score organization in Prov-GigaPath embedding space without imposing the full diagnostic sequence during graph-distance computation. Without assuming a strict linear progression for intermediate lesions or individual ROIs, the analysis supports a trajectory-like class-level organization across breast lesion categories.

### Pairwise Energy distance and RBF-MMD² reveal graded distributional separation

We next tested whether the graph-distance organization was supported by a complementary, order-free comparison of class-specific embedding distributions. For each split, we computed pairwise Energy distances between diagnostic categories and used RBF-MMD² as a complementary sensitivity analysis. In practical terms, these metrics quantify how dissimilar the overall embedding distributions of two lesion categories are. These analyses used class labels to define the groups being compared, but did not impose an ordinal class sequence or specify which class pairs should be adjacent.

Energy-distance matrices showed a broad pattern in which widely separated diagnostic categories, including N and IC, had larger distributional distances, whereas several diagnostically proximate category pairs had smaller distances. This broad organization was observed across the training, validation, and test splits (**Fig. 4A–C**). RBF-MMD² analysis at the training-derived reference bandwidth in the training split showed a similar pairwise organization, and the broad pairwise organization was retained across alternative RBF bandwidths and the three data splits, corroborating that the observed distance structure was not specific to a single distributional metric or RBF bandwidth (**Fig. 4D**). To assess within-patient dependence and quantify patient-level resampling variability, we performed a global patient-block bootstrap in which patients, rather than individual ROIs, were resampled as blocks and representative pairwise Energy distances were recomputed. Patient-block bootstrap analyses further supported the stability of representative pairwise Energy distance estimates, including larger separation for distant class pairs such as N–IC compared with diagnostically close category pairs (**Fig. 4E**). Thus, order-free Energy distance and RBF-MMD² analyses together support a broadly graded distributional organization among BRACS lesion categories in the Prov-GigaPath morphology space.

**Fig. 4.**
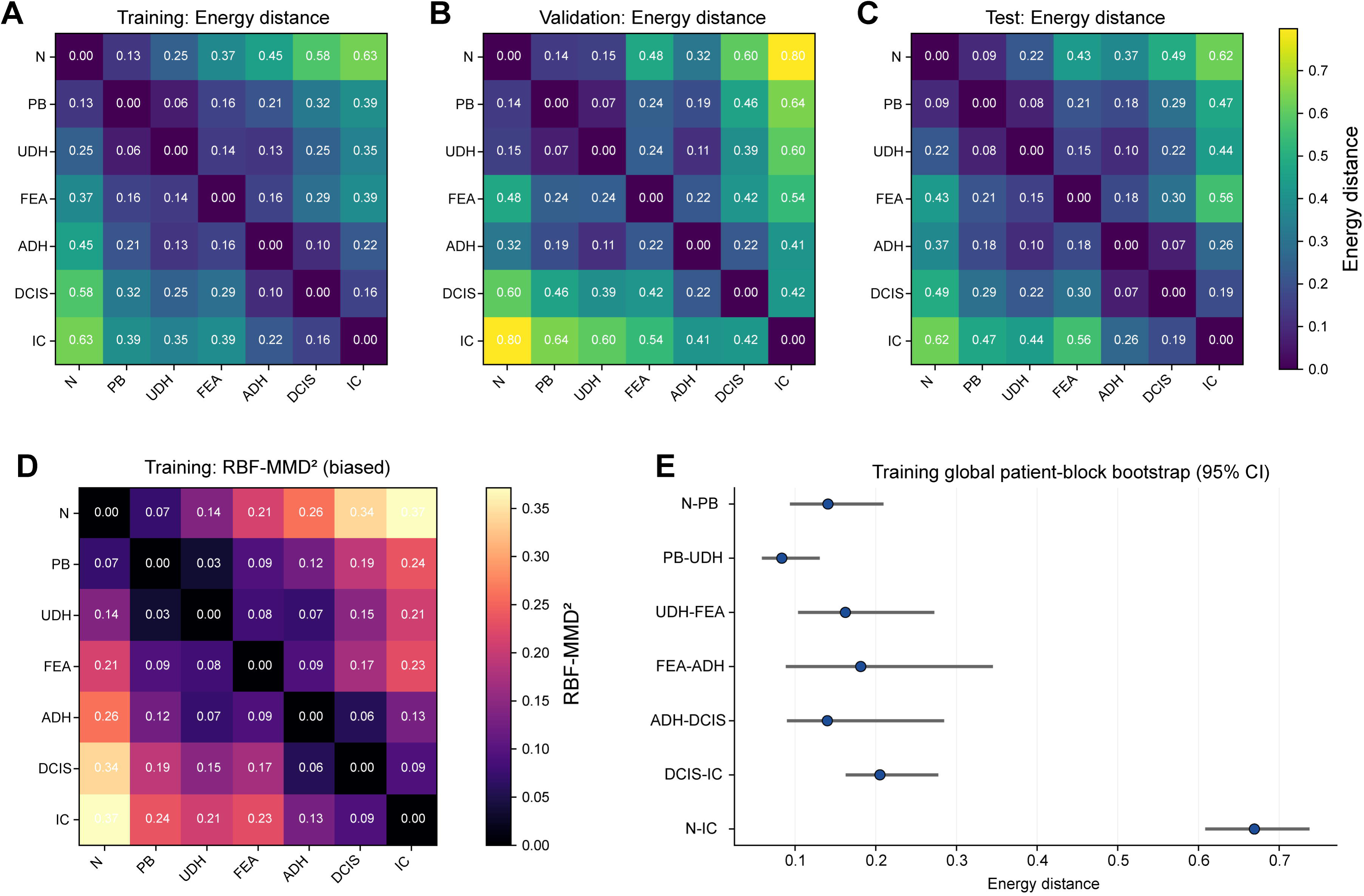
Order-free pairwise distributional distances across BRACS lesion categories. (A–C) Pairwise biased V-statistic Energy-distance matrices for the training, validation, and test splits, respectively, calculated in the primary train-fitted PCA representation. Diagnostic labels defined the distributions being compared, but no ordinal sequence or adjacency structure was used in the distance calculation. Lower values indicate more similar class-specific ROI embedding distributions, whereas higher values indicate greater distributional separation. The diagonal was set to zero, and a common color scale was used across panels A–C. **(D)** Training-split biased RBF-MMD² matrix at the train-defined reference bandwidth. **(E)** Representative training-split global patient-block bootstrap medians and 95% confidence intervals for selected class pairs. The displayed pairs include diagnostically close contrasts and the broad N-IC contrast; pair selection does not enter the Energy-distance calculation.

### Embedding-defined morphology space captures within-DCIS heterogeneity associated with annotation-derived invasive-carcinoma context

We next leveraged the embeddings to dissect biologically and pathologically derived questions. DCIS associated with synchronous invasive carcinoma has been reported to show more aggressive clinicopathologic features and/or molecular features closer to invasive carcinoma than pure DCIS(10–12). We asked whether the embedding-defined morphology space captured within-DCIS heterogeneity related to invasive-carcinoma context.

As a pathology-informed post hoc analysis, we tested whether the Normal-rooted morphology score could reveal within-category heterogeneity among DCIS ROIs according to BRACS annotation-derived IC co-observation. Given that BRACS provides ROI/WSI annotations rather than component-level spatial linkage or longitudinal clinical confirmation, we defined the primary context by same-WSI IC co-observation and did not interpret the no-observed-IC group as clinically and pathologically confirmed pure DCIS.

After the patient-level split audit and exclusion of the overlapping patient, the analysis included 780 DCIS ROIs from 87 WSIs and 49 patients. The primary independent-patient comparison excluded five mixed-context patients, yielding 26 patients with no observed same-WSI IC and 18 patients with observed same-WSI IC (**Fig. 5A**). In the train-defined PC1–PC2 display, WSI-balanced patient-level DCIS points from the observed same-WSI IC group showed a modest visual shift away from the N-associated region and toward the IC-associated side of the space, without forming discrete clusters (**Fig. 5B**).

**Fig. 5.**
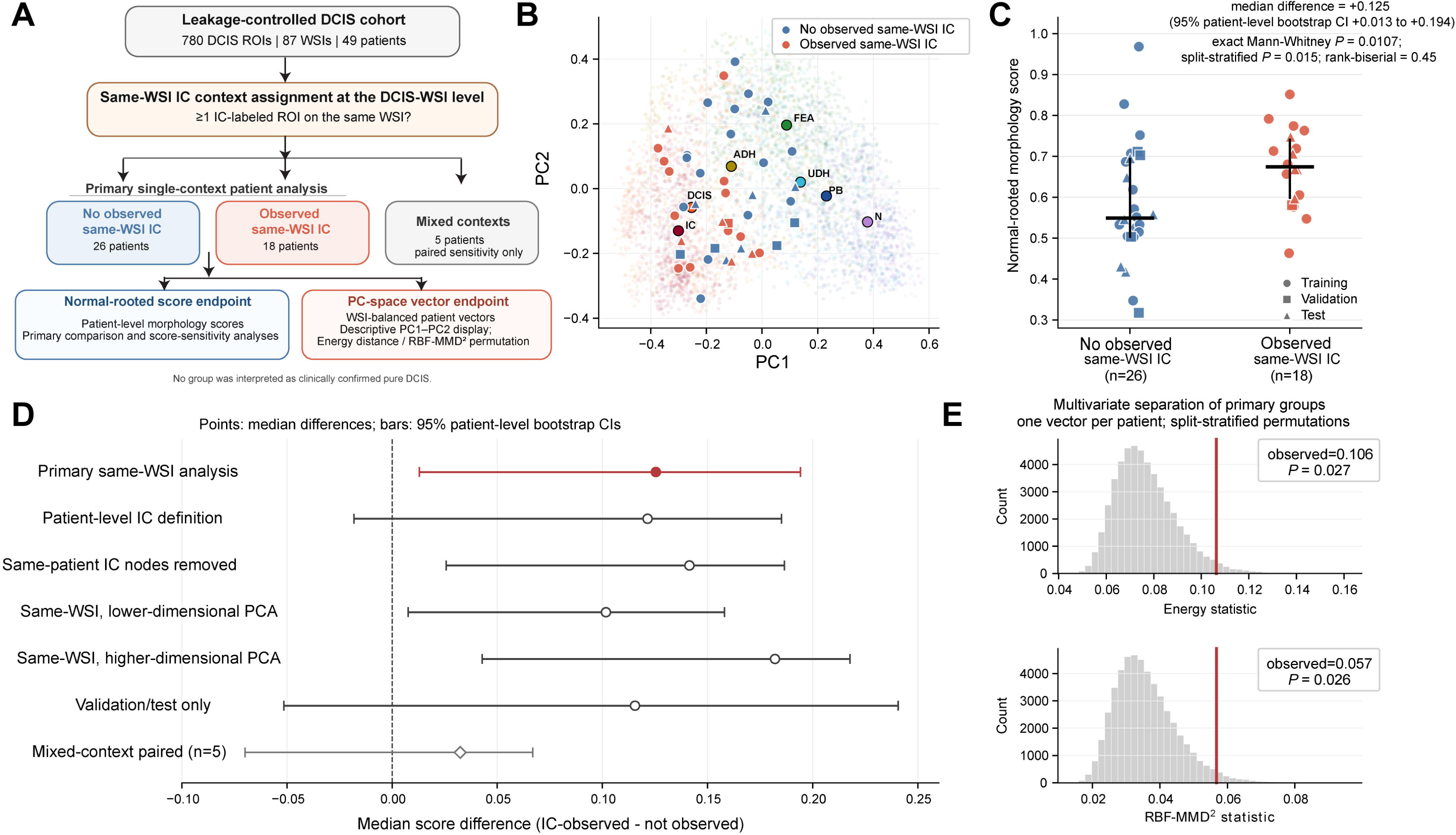
DCIS morphology-space heterogeneity according to annotation-derived invasive-carcinoma context. **(A)** Overview of the DCIS context analysis. Same-WSI IC co-observation was defined by the presence of at least one IC-labeled ROI on the same WSI. The primary comparison excluded five mixed-context patients and compared independent patient groups after WSI-to-patient aggregation. Patients with DCIS WSIs exclusively in one context were assigned to the primary no observed same-WSI IC or observed same-WSI IC groups, whereas patients contributing DCIS WSIs in both contexts were classified as mixed-context patients. The broader patient-level IC definition was used only as a sensitivity analysis and was based on whether any IC annotation was observed in the available BRACS material. Both endpoints were analyzed at the patient level, accounting for the WSI structure. **(B)** Descriptive display of primary-analysis DCIS patient vectors, from which mixed-context patients were excluded, in the train-defined PC1–PC2 space. Faint points show retained BRACS ROIs for orientation, labeled points show class centroids and colored points show one aggregated DCIS vector per patient. This two-dimensional display was not used for inference. **(C)** Primary patient-level comparison of Normal-rooted kNN-Dijkstra morphology scores. Each point represents one patient, with marker shapes indicating the training (circle), validation (square), or test (triangle) split. **(D)** Sensitivity analyses for the score difference across context definitions, graph-reference handling, PCA dimensionality and split restriction. Points show median score differences and horizontal bars show 95% percentile bootstrap confidence intervals (CIs) based on patient-level resampling. **(E)** Patient-level multivariate distributional comparison using Energy distance and RBF-MMD². Histograms show split-stratified label-permutation null distributions; vertical lines indicate observed statistics. These exploratory analyses assess annotation-associated within-DCIS morphology-space heterogeneity and do not establish spatial adjacency, clonal relatedness, temporal progression, or invasive potential; the no-observed-IC group was not interpreted as clinically confirmed pure DCIS.

In the primary patient-level analysis, DCIS from the observed same-WSI IC group had higher Normal-rooted morphology scores than DCIS from the no-observed same-WSI IC group (median difference, +0.125; 95% patient-level bootstrap confidence interval (CI), +0.013 to +0.194; exact Mann–Whitney *P* = 0.0107; split-stratified permutation *P* = 0.015; **Fig. 5C**). The direction of the score difference was retained under a broader patient-level IC-annotation definition, after removal of same-patient IC nodes from the graph reference, and in lower- and higher-dimensional PCA analyses, whereas the validation/test-only estimate was directionally similar but not statistically conclusive. The five mixed-context patients did not provide independent paired confirmation (**Fig. 5D**).

Complementing the one-dimensional score analysis, we represented each patient’s DCIS morphology as a single WSI-balanced point in the primary PCA space and tested whether the two primary patient groups, defined by the absence or presence of observed same-WSI IC, were more separated than expected after random reassignment of the context labels within each data split. Only 2.7% and 2.6% of the split-stratified label permutations yielded Energy-distance and RBF-MMD² values, respectively, at least as large as the observed values (Energy distance = 0.106, nominal P = 0.027; RBF-MMD² = 0.057, nominal P = 0.026; **Fig. 5E**). Comparable results were obtained in lower- and higher-dimensional PCA spaces (both metrics, nominal P = 0.027–0.036).

These findings suggest that the embedding-defined morphology space can capture within-DCIS heterogeneity related to annotation-derived IC co-observation.

## Discussion

In this study, we identified reproducible morphology-space organization across BRACS lesion categories from frozen Prov-GigaPath ROI embeddings, using a leakage-controlled, patient-aware framework. This organization was supported by train-defined PCA visualization, Normal-rooted kNN-Dijkstra graph distances and order-free pairwise distributional distances and was retained across validation/test projection and patient-aware sensitivity analyses. Together, these results show that breast lesion classes are organized along a reproducible embedding-space morphology axis from normal/benign to malignant categories, supporting trajectory-like relationships in embedding-defined morphology space.

Our study extends the application of pathology foundation-model embeddings beyond discrete diagnostic classification by mapping breast lesion categories within a shared morphology state space. These findings are consistent with a broader shift in computational pathology from task-specific diagnostic classification toward reusable representation learning, biomarker prediction, prognosis, retrieval, and morphology-state modeling(1, 23). Recent advances in pathology foundation models have made it possible to use frozen or lightly adapted embeddings as transferable morphology representations, rather than training a separate classifier for each diagnostic endpoint(1, 15, 24). Recent computational studies have further suggested that histopathological morphology can be modeled as a continuous biological state rather than only as a categorical diagnostic endpoint (25, 26). However, whether distributional embeddings can quantitatively organize the clinically and pathologically challenging spectrum across diagnostic categories, such as benign, atypical, in situ, and invasive breast lesions, remains unclear. Our study addresses this gap by showing that frozen pathology foundation-model embeddings can organize histopathology ROIs spanning multiple breast lesion categories within a reproducible embedding-space morphology state space, with broad positioning from normal/benign to malignant lesions, without training a tumor-specific classifier or imposing the full diagnostic sequence.

Furthermore, the post hoc DCIS context analysis suggested that foundation-model embeddings can capture clinicopathologically relevant heterogeneity within a conventional diagnostic category. Our analysis showed that DCIS from the observed same-WSI IC group had higher Normal-rooted morphology scores than DCIS from the no-observed same-WSI IC group, while complementary patient-level multivariate analyses showed context-associated distributional separation. Prior clinicopathologic and molecular studies reported that DCIS associated with synchronous invasive carcinoma can show more aggressive pathologic features and molecular profiles closer to invasive carcinoma than pure DCIS(10–12). Importantly, the Normal-rooted score difference and multivariate distributional separation were identified without using molecular information in the analysis, and the score difference was not attributable solely to same-patient IC nodes in the training graph. The concordance between the image-derived morphology-space findings and previously reported clinicopathologic and molecular differences supports the possibility that foundation-model embeddings may add clinicopathologically relevant granularity within diagnostic labels, rather than only separating predefined categories.

Several limitations should be considered. First, because this study used cross-sectional BRACS ROIs rather than longitudinally sampled lesions, the inferred morphology-space organization should be interpreted as a trajectory-like structure, not as direct evidence of temporal progression, clonal evolution, or causal biological transition. Second, although the kNN-Dijkstra analysis did not impose the full diagnostic sequence, it used the Normal-class training ROI as the graph root, and the Energy-distance and RBF-MMD² analyses used diagnostic labels to define the compared distributions. Thus, the framework is best viewed as a patient-aware morphology-space mapping approach rather than a fully label-free trajectory discovery method. Third, the DCIS context analysis relied on annotation-derived same-WSI IC co-observation rather than component-level spatial linkage or longitudinal clinical confirmation; therefore, these findings should be interpreted as exploratory evidence of morphology-space heterogeneity within DCIS. Fourth, this study used a single public ROI-level dataset, a single foundation model, and a fixed training reference space. Nevertheless, validation and test projections retained broad class-rank concordance, including after restriction to in-reference ROIs, supporting the robustness of the overall morphology-space organization. BRACS was well suited to this proof-of-concept analysis because its seven consensus-annotated categories span normal tissue, benign and atypical lesions, DCIS and IC(14). The inclusion of UDH, FEA and ADH enabled evaluation of intermediate morphology states, rather than only separation between benign and malignant lesions. Future studies in independent whole-slide cohorts with component-level pathology, receptor status, histologic/nuclear grade, molecular profiles, treatment, and outcome annotations should extend this framework across additional foundation models and define the biological and clinical significance of morphology-space positioning.

## Conclusions

Taken together, our approach provides a framework for mapping breast lesion histopathology within a morphologic state space, enabling tissue heterogeneity and histologic severity to be represented as continuous positioning rather than classification alone. By leveraging foundation-model embeddings and morphology-space mapping, this framework may help uncover latent pathobiological information embedded in routine histopathology. More broadly, this framework may extend the clinical utility of routine histopathology by helping to guide targeted molecular follow-up, inform more individualized risk assessment, and support cost-efficient clinicopathological decision-making.

## Declarations

### Ethics approval and consent to participate

This study was a secondary analysis of publicly available, de-identified data from the BRACS dataset using the publicly available Prov-GigaPath model (see Availability of data and materials). No new human participants were recruited, and no new human tissue samples or identifiable personal data were collected for this study. Therefore, ethics approval and consent to participate were not applicable to this study.

### Consent for publication

Not applicable.

### Availability of data and materials

The dataset and model supporting the conclusions of this article are available in the BRACS repository, https://www.bracs.icar.cnr.it/, and Prov-GigaPath, https://github.com/prov-gigapath/prov-gigapath.git, respectively.

### Authors’ contributions

Y.M. conceptualized the study, performed data analysis, prepared the figures, wrote the original manuscript, and supervised the study. E.T. contributed to data analysis. T.S. supported pathological interpretation and supervised the study. All authors read and approved the final manuscript.

## Acknowledgements

We thank members of the Department of Anatomic Pathology, Tohoku University Graduate School of Medicine, SiRIUS Institute of Medical Research, Tohoku University, AI Lab and the Department of Pathology, Tohoku University Hospital for their support.

## Abbreviations

ADH: Atypical ductal hyperplasia
BRACS: BReAst Carcinoma Subtyping dataset
CI: Confidence interval
DCIS: Ductal carcinoma in situ
FEA: Flat epithelial atypia
H&E: Hematoxylin and eosin
IC: Invasive carcinoma
kNN: k-nearest neighbor
MMD²: Squared maximum mean discrepancy
N: Normal tissue
NAC: Neoadjuvant chemotherapy
PB: Pathological benign lesion
PC: Principal component
PCA: Principal component analysis
QC: Quality control
RBF: Radial basis function
RBF-MMD²: Squared maximum mean discrepancy with a radial-basis-function kernel
ROI: Region of interest
UDH: Usual ductal hyperplasia
WSI: Whole-slide image

